# Latitudinal gradient in species diversity provides high niche opportunities for a range-expanding phytophagous insect

**DOI:** 10.1101/2022.02.07.479421

**Authors:** Dylan G. Jones, Julia Kobelt, Jenna M. Ross, Thomas H.Q. Powell, Kirsten M. Prior

## Abstract

1. When species undergo poleward range expansions in response to anthropogenic change, they likely encounter less diverse communities in new locations. If low diversity communities provide weak biotic interactions, such as reduced competition for resources or predation, range-expanding species may experience ‘high niche opportunities.’
2. Here, we uncover if oak gall wasp communities experience a latitudinal diversity gradient (LDG) and weaker interactions at the poles that might create high niche opportunities for a range-expanding community member.
3. We performed systematic surveys of oak gall wasps on a dominant oak, *Quercus garryana*, throughout most of its range, from northern California to Vancouver Island, British Columbia. On 540 trees at 18 sites, we identified 23 species in three guilds (leaf detachable, leaf integral, and stem galls). We performed regressions between oak gall wasp diversity, latitude, and other abiotic and habitat factors to reveal if cynipid communities follow an LDG. To uncover patterns in local interactions, we performed partial correlations on oak gall wasps co-occurring on trees within regions. Finally, we performed regressions between abundances of co-occurring gall wasps on trees to reveal potential interactions.
4. *Q. garryana-gall* wasp communities followed an LDG, with lower diversity at higher latitudes, particularly with a loss of detachable leaf gall species. Detachable leaf gall wasps, including the range-expanding species, co-occurred most on trees and had weak interactions in the northern region. Abundances of the range-expander and detachable and integral leaf galls co-occurring on trees were negatively related, suggesting antagonistic interactions. Overall, we found that LDGs create communities with weaker antagonistic interactions at the poles that might facilitate ecological release in a range-expanding community member.
5. Here, we uncover how regional and local scale patterns and processes create high niche opportunities for a range-expanding community member. This research provides insight into how biogeographical patterns in communities and species interactions influence the outcome of range expansions. Given the ubiquity of LDGs, these outcomes might be expected in other phytophagous insect communities.

## Introduction

Anthropogenic change is causing species to shift their ranges, including to higher latitudes and elevations (Parmesan and Yohe 2003). Species experience range shifts at different rates due to variation in thermal tolerances and dispersal capabilities (Parmesan and Yohe 2003, Hellmann et al. 2012). As a result, range-expanding species may leave interacting species behind and form novel interactions as they infiltrate new species pools (Gilman et al. 2010, Hellmann et al. 2012). Depending on the net effects of lost or gained antagonistic and facilitative interactions, range-expanding species may experience ‘high niche opportunities’ (*sensu* Shea and Chesson 2002) resulting from increased resource availability or reduced predation that results in higher performance in new locations ‘ecological release’ or low niche opportunities or ‘biotic resistance’ leading to lower performance (Shea and Chesson 2002, Gilman et al. 2010).

One of the most ubiquitous patterns in nature is that species diversity declines towards the poles or higher elevations (Latitudinal Diversity Gradient, LDG) (Gaston 2000, Willig et al. 2003, Hillebrand 2004, Fitt and Lancaster 2017, Zvereva and Kozlov 2021). This pattern has been observed across many taxa, including phytophagous insects and parasitoids (Cuevas-Reyes et al. 2003, Willig et al. 2003, Kerr et al. 2016, Galiana et al. 2019). LDGs are often linked to latitudinal gradients in biotic interactions, with weaker interactions towards the poles (Pennings et al. 2009; Schemske et al. 2009, Zvereva and Kozlov 2021, except see Moles et al. 2011). Low-diversity communities are predicted to have high niche opportunities if there are weak interactions among competitors or with predators (Shea and Chesson 2002, Willing et al. 2003, Schemske et al. 2009, Gilman et al. 2010). As a result, poleward communities may provide weak biotic resistance, making high latitude communities susceptible to colonization by range-expanding species and invaders (Gilman et al. 2010, Wallingford et al. 2020). Understanding if LDGs provide high niche opportunities is important to uncover and predict potential outcomes of range expansions.

While LDGs result from regional-scale processes, uncovering whether gradients provide high niche opportunities requires linking regional and local scale patterns and processes (Shea and Chesson 2002, Fitt and Lancaster 2017, Wallingford et al. 2020). Patterns in species diversity occur through a series of ‘filtering’ processes. Across a geographic range, regional species pools are determined by large scale processes, including gradients in abiotic conditions and dispersal after historic geographical events (Gaston 2000, Willig et al. 2003, Bello et al. 2013, Deák et al. 2018). These factors limit species distributions generating variation in regional species pools across latitude. Local species pools are determined by a combination of processes, including dispersal from regional pools, suitability of abiotic conditions, and biotic interactions (Chase 2011, Cornell and Harrison 2014). Response to high niche opportunities depends on release from biotic interactions at local scales (Shea and Chesson 2002, Bello et al. 2013).

Phytophagous insects do not often compete directly, but more commonly interact indirectly mediated by direct interactions with host plants or enemies (Veen et al. 2006, Kaplan and Denno 2007, Prior and Hellmann 2010, Holt and Bonsall 2017). Lower phytophagous insect diversity at the poles may create high niche opportunities on host plants by relaxing plant-mediated competition, leading to susceptible host plants with more available resources or low defenses (Woods et al. 2012, Wieski and Pennings 2014). Release from ‘apparent competition’ (competition via shared enemies) at the poles could result from lower abundances of host or enemy diversity, or if enemies fail to effectively switch to attacking novel hosts (Shea and Chesson 2002, Menendez et al. 2008). There is evidence for patterns of weaker biotic interactions between phytophagous insects and host plants at the poles, with studies finding lower host plant damage or defenses or higher foliar nutrients (Pennings et al. 2009, Wieski and Pennings 2014). Previous studies of range-expanding phytophagous insects have also found evidence for weaker apparent competition, resulting from lower host and enemy diversity, and from enemies failing to effectively attack novel hosts (Schönrogge et al. 2000, Menendez et al. 2008, Prior and Hellmann 2013).

Here we leverage a poleward range expansion of a phytophagous oak gall wasp, *Neuroterus saltatorius* (Hymenoptera: Cynipidae) (cynipid) that occurs a dominant oak in western North America, *Quercus garryana*. *Q. garryana* hosts a community of cynipid wasps that co-occur with *N. saltatorius* that are attacked by a community of parasitoid wasps (Smith 1995, Prior and Hellmann 2013). The native range of *N. saltatorius* on *Q. garryana* spans from California to Puget Sound, Washington (Russo 2006, Prior and Hulcr 2017). This species recently expanded its range poleward to Vancouver Island, BC and now occurs at the most northern locations of *Q. garryana* (Duncan 1997, Prior and Hellmann 2013). In its expanded range, it occurs at higher abundances on a higher proportion of trees compared to its native range (Prior and Hellmann 2013).

The objective of this study is to test whether the *Q. garryana-cynipid* community follows an LDG, and if lower diversity at the poles provides weak interactions for the range-expanding species. To this end, we measure cynipid diversity throughout the range of *Q. garryana* to uncover if diversity follows an LDG, and if other factors contribute to patterns in regional diversity. Next, we examine how regional species pools influence local patterns of cynipids that co-occur and putatively interact on individual host plants. Finally, we uncover if co-occurring cynipids interact with *N. saltatorius* as evidence that release from local antagonistic interactions contribute to ecological release. We predict that *Q. garryana-cynipid* communities follow an LDG and that regional species pools create variation in local niche opportunities, with higher niche opportunities at the poles via reduced interactions on host plants.

## Methods and Materials

### Study system and sites

*Quercus garryana* var. *garryana* Douglas ex. Hook (Fagaceae), Garry/Oregon white oak, ranges from central California to southwestern BC. *Q. garryana* occurs in semi-arid woodlands or savannas and is often the predominant overstory vegetation (Stein 1990). *Q. garryana* is the only oak species from central Oregon to BC but overlaps with several oak species in the southern portion of its range (Stein 1990). Oak ecosystems occur in rain shadows of coastal mountains, and their distribution becomes patchier (surrounded by *Pseudotsuga menziesii* forests) with increasing latitude, due to decreased suitable abiotic conditions (Stein 1990, Vellend et al. 2008).

Cynipid wasps are a specialized group of phytophagous insects that deposit their eggs in living plant tissue of Fagaceae (beech and oaks), eliciting the formation of a gall structure that houses larvae while it feeds on plant tissue (Stone et al. 2002, Hayward and Stone 2005). Cynipids often have two generations; a gamic (sexual) generation and an agamic (asexual) generation that form distinct galls. Cynipid galls vary significantly in morphology among species and among generations and occur on various plant tissues (Stone et al. 2002, Hayward and Stone 2005, Russo 2006). There are ~700 described oak cynipid species in North America, with western oak ecosystems being a hotspot of diversity (Weld 1957, Burks 1979, Russo 2006). Cynipid wasps are attacked by a suite of parasitoids wasps, largely in the Superfamily Chalcidoidea, many of which are polyphagous within cynipids, but generally specialized on cynipids (Hayward and Stone 2005, Bailey et al. 2009, Askew et al. 2013, Forbes et al. 2016).

*Neuroterus saltatorius* (Edwards), the jumping gall wasp, is one of several cynipid wasps on *Q. garryana* in western North America (Russo 2006, Prior and Hulcr 2017). *N. saltatorius’* s native range on *Q. garryana* occurs from northern California to Puget Sound, Washington; and recently expanded its range to the northernmost extent of *Q. garryana* in BC (Duncan 1997). This species is absent in historical records (Evans 1985), and its first appearance and spread in BC was tracked by the Canadian Forest Service starting in 1983 around the city of Victoria (Duncan 1997), and Prior and Hellmann (2013) noted that it occurred at the northern most intact *Q. garryana* site in BC on Hornby Island by 2007.

*N. saltatorius* forms small galls (1-2 mm), with clustered integral leaf galls produced by the gamic spring generation and detachable leaf galls formed by the summer agamic generation (Smith 1995). The detachable agamic galls drop from the leaves in mid-late summer where the galls remain in the leaf litter for the winter until the following spring (Smith 1995). The agamic generation occurs at high abundance on *Q. garryana* in its expanded range, with more infested trees. There are infested trees in the native range, but they are usually isolated (Prior and Hellmann 2013). High abundance causes foliar scorching with putative effects on oaks and documented effects on native oak insects (Duncan 1997, MacDougall 2010, Prior and Hellmann 2010).

We selected *Q. garryana* study sites based on oak dominance, patch size, and accessibility. We predetermined six regions based on distribution and logistics, with three sites in each region. Sites in regions were separated on average 44.67 km, with closest sites being 3 km from one another. All oak patches were >1.5ha (Fig. 1; Appendix S1:Table S1). Sites were managed by various landowners and varied in management practices. None of the sites had visible signs of recent burning, and we avoided choosing sites with known history of recent burns.

**Figure 1:**
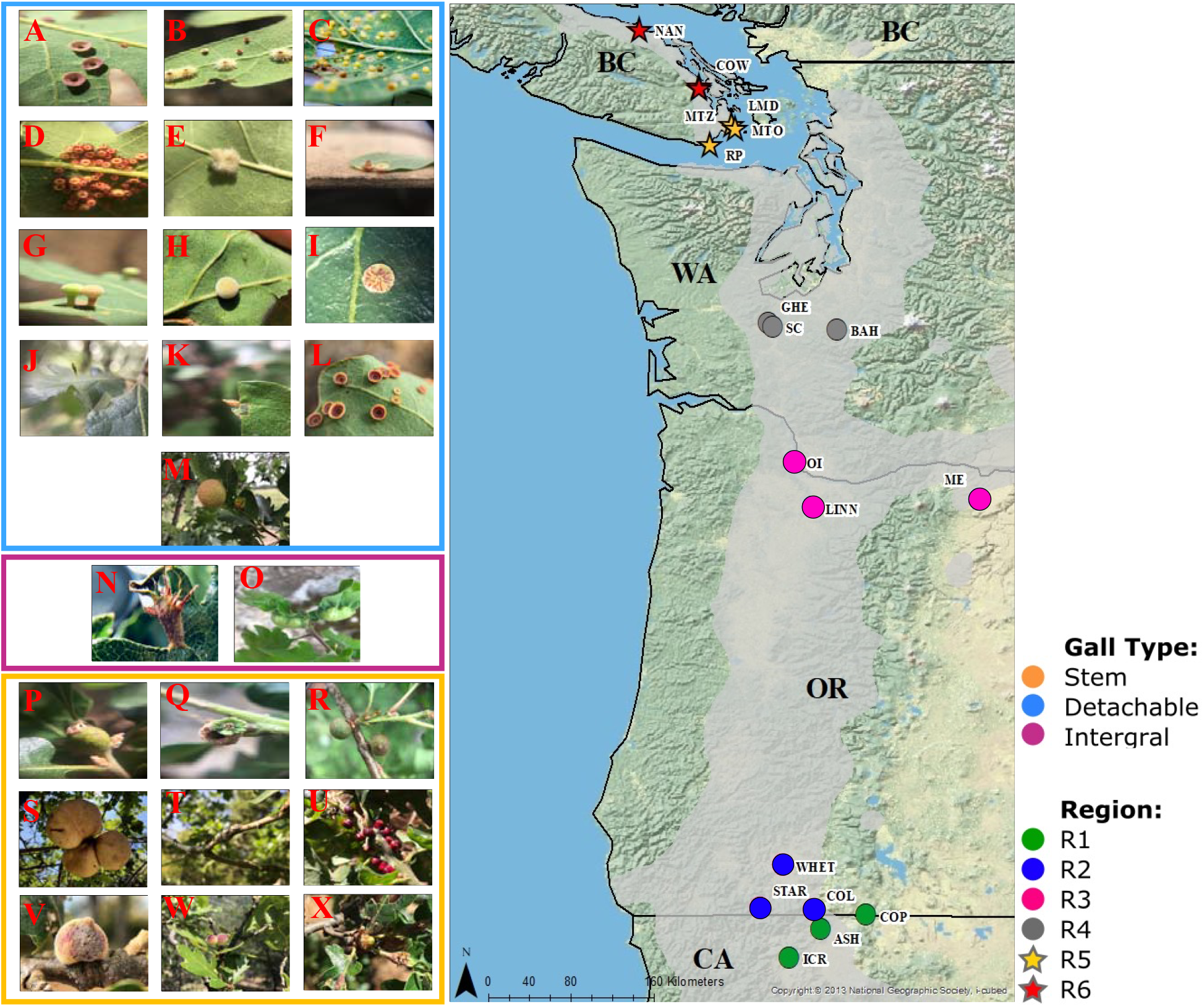
Cynipid morphotypes found on *Quercus garryana*: Row 1) *Andricus gigas* (*AG*), *A. stellaris (AS), Neuroterus saltatorius* (agamic, *NSA*), Row 2) *A. tubifaciens* (TT), *A. confertus* (*ACF*), *A. kingi* (agamic, *AKA*), Row 3) *Xanthoteras teres* (*XT*), *A. fullawayi* (*AF*), *A. parmula* (AP), Row 4) *A. pedicellatum* (*DP*), *A. kingi* (gamic, *AK*), Unknown *sp*. (pink bowtie, *PB*), Row 5) *Besbicus mirabilis* (*BMI*), *A. opertus* (*AO*), *N. washingtonensis* (*NW*), Row 6) *A coortus* (*AC*), *Unknown sp*. (acorn cap, *ACG*), *Burnettweldia washingtonensis* (*DW*), Row 7) *A. quercuscalifornicus* (*AQ*), *A. chrysolepidicola* (*ACH*), *Disholcaspis mellifica* (*DME*), Row 8) *D. canescens* (*DC*), *D. mamillana/simulata* (*DM/DS*), *Unknown* sp. (acorn gall, *ACO*). Morphotypes colored by gall type guild (blue = detachable leaf, purple = integral leaf, orange = stem). Y) map showing range of *Quercus garryana* (grey) and study sites grouped into six regions represented by different colors, circles represent sites in the native range, stars expanded range (see Table S1 for site acronyms).

### Sampling cynipid communities

We employed a systematic, uniform-effort sampling regime to produce comparable abundance and community composition data across sites. We visited each site three times (sampling periods) to capture temporal changes in cynipid communities that coincided with the development time on trees of the agamic generation of *N. saltatorius* (Smith 1995). We focused on the agamic generation as it is the more abundant and damaging generation (Duncan 1997, Prior and Hellmann 2010). We started sampling sites when *N. saltatorius* agamic galls started to develop, with the first period beginning on May 21, 2019 in the southernmost region (R1) and continued in order of increasing latitude. Sampling of the northern regions (R4-R6) began on June 5^th^, 2019 (Appendix S1:Table S1). We sampled one site per day, and regions were sampled in the same order throughout the season, with an average of 20 days between site visits.

During each visit, we haphazardly sampled 10 trees without replacement, for a total of 30 trees per site. Trees were at least 10 m apart from previously sampled trees, had enough reachable limbs to sample (up to 8 feet, standing on a 6-foot ladder), and contained at least one cynipid when searching during the first 5 minutes of sampling. We standardized effort by sampling 10 branches, where we exhaustively searched for galls on 10 leaf clusters per branch and on stems on approximately 1 m. Galls occur on other structures, such as buds; however, the leaves were already flushed out by our first sampling date, and it is possible that cryptic species of gall wasps were not recorded. We did not search in the higher canopy of the trees; previous studies found no variability in cynipid richness between the different stratifications of the canopy (Eliason and Potter 2001).

Gall morphotypes were identified via gall morphology (Weld 1957, Russo 2006). Some gall species were difficult to identify via their gall structure alone without destroying the galls, and in one case we lumped species: *Disholcaspis mamillana* and *D. simulata.* (Fig. 1, Appendix S1:Table S2). Total gall abundance for each cynipid morphotype was recorded for each tree. We put galls into three gall-type guilds: integral leaf galls, detachable leaf galls, and non-leaf galls (that included mostly stem galls, but also petiole and acorn galls; *hereafter* ‘stem galls’) (Russo 2006). *N. saltatorius* agamic generation is a detachable leaf gall (Russo 2006).

### Measuring abiotic and habitat variables

On the first survey day at each site, we hung an iButton temperature logger (1-Wire, Thermochron) on one tree to collect temperature hourly. Loggers were kept up until the final sampling date. For each survey tree, we measured soil temperature and moisture at the base of the tree. We measured tree height (m) and diameter at breast height (DBH) and created a composite ‘tree size’ metric (Appendix S1:Supplementary Methods). For each site, we collected BioClim variables with a spatial resolution of ~340 km^2^ from WorldClim 1970-2000 data: temperature and precipitation (annual mean, max., min., and seasonality) (Fick 2017) and created separate composite temperature and precipitation variables using Principal component analysis (PCA) (Appendix S1: Supplementary Methods; Fig.S1). We chose these variables as they contribute to patterns in insect species distributions (Gon□alves-Souza et al. 2014, Moreira et al. 2015). We also measured survey area, oak patch size, and determined broad habitat and land use categories surrounding survey areas using aerial images (Appendix S1:Supplementary Methods).

### Determinants of regional patterns in cynipid diversity

To assess if predetermined regions reflected turnover in regional diversity, we partitioned cynipid richness at several spatial scales (within (α) and among (ß) regions, sites, trees) by performing additive partitioning using the program PARTITION (Veech and Crist, 2009). We found significant turnover in ß-richness among regions but not among sites in regions (Appendix S1:Table S4; Fig.S3). Thus, we defined the regional species pools in pre-determined regions and pooled trees among three study sites in our local scale analysis (see below, 90 trees per region).

We calculated cynipid morphotype richness for each survey period at each site. We created rarefaction curves that justified our use of raw richness for subsequent analyses (Appendix S1:Fig. S4). To describe cynipid community composition, we performed a canonical analysis (CA) that is a suitable ordination approach to assess species composition over gradients (Legendre 2008) (vegan package in R, Oksanen et al. 2013). We used a matrix of cynipid morphotypes abundances observed during surveys at sites and used CA1 and CA2 as metrics of community composition for models (see below). We also created a site level CA biplot (Fig. 2).

**Figure 2:**
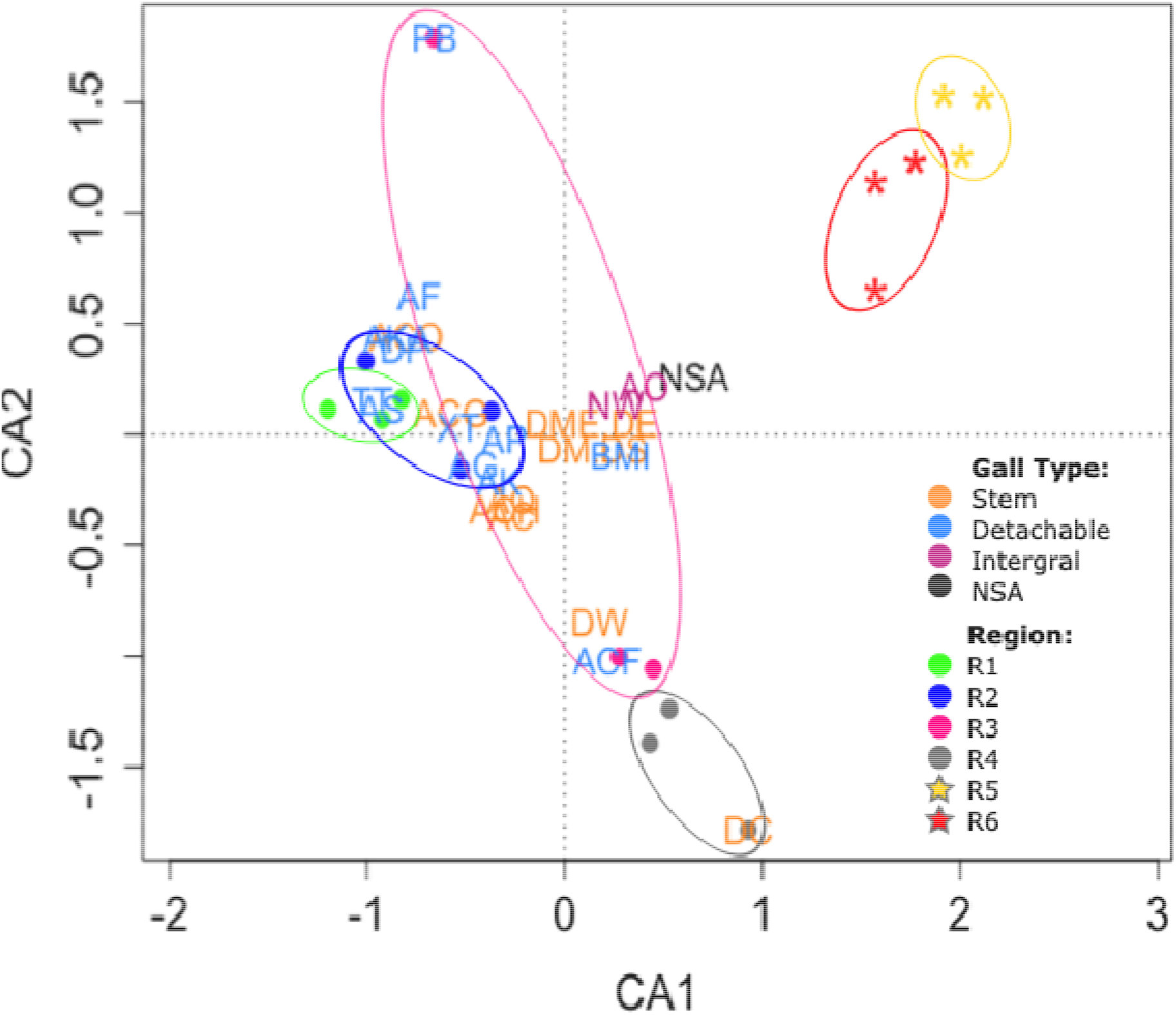
Canonical Analysis (CA) biplot of cynipid morphotype composition at sites across regions. Circles denote sites in the native range, and stars in the expanded range. Ellipses encompass sites within regions. Gall morphotype and site abbreviations Fig. 1, colored gall acronyms represent gall type guild (blue = detachable leaf, purple = integral leaf, orange = stem) (Tables S1, S2).

To determine if patterns in cynipid richness, composition, and abundance follow an LDG and if abiotic and habitat factors contribute to diversity patterns, we ran linear (LMM) or generalized linear mixed effect models (GLMM) with multiple predictor variables as fixed effects (Appendix S1:Table S3), and survey date (Julian date) and region as random effects. Since our main objective includes evaluating the LDG, we included latitude in all models. Sites also varied in elevation which we also included. We included other abiotic and habitat variables that may contribute to patterns in cynipid diversity, some that correlate with latitude, including climate factors (PC_temp_, PC_precip_, soil moisture) and habitat factors (oak patch size, PCh_abitat types_) (Appendix S1:Fig.S1, Table S3). First, we ran mixed effect models with single predictor variables and calculated conservative Akaike information criterion (AICc) scores for each model. We then built models by creating multi-factor models using a forward stepwise approach starting with variables with the lowest AICc scores, and then building to add in all variables. We compared and retained models with a ΔAICc < 2 (Hill et al. 2017). (see Appendix S1: Supplementary methods for details).

### Local patterns in cynipid interactions

To assess potential competitive interactions and variation in niche opportunities among regions, we determined cynipid morphotype associations on trees, as sedentary phytophagous insects most strongly interact on individual host plants (Kaplan and Denno 2007). Given that diversity is relatively homogenous among sites within regions (Appendix S1:Fig. S3, Table S4), we pooled sites, using 90 trees per region. We used ggcorrplot to perform partial correlation analyses in R using presence/absence data of gall morphotypes on trees (Kassambara 2019). Correlations were categorized as weak (*R* = 0.25-0.49), moderate (*R* = 0.50-0.74.5), or strong associations (*R* = 0.75-1.0). Correlation values less than 0.25 were categorized as no association. For correlations between morphotypes > 0.25, we performed Spearman’s Rank Order Correlation tests to determine if correlations were significant between morphotype pairs in each region. igraph in R was used to create association networks (Csardi 2020). We performed canonical correspondence analysis (CCA) with presence/absence data with several predictor variables to uncover if site variables influenced tree community composition (Appendix S1:Fig. S7).

Finally, to assess if associations on trees reflect putative interactions (with the focal species), we performed LMs between the abundance of other cynipid morphotypes and the abundance of *N. saltatorius* using trees where they co-occur. We performed separate analyses for each gall-type guild (detachable, integral, stem) as we predict overlapping gall types might more strongly interact. We scaled cynipid abundances by dividing the number of galls on trees for each group (*N. saltatorius*, detachable, integral, and stem) by the total number of galls at a site to make comparisons among sites with different abundances. We included other gall abundance, region, and the interaction as fixed effects. We performed arcsine square root transformations on relative gall abundances. Significant negative relationships provide evidence of antagonistic interactions between other cynipids and *N. saltatorius*.

## Results

### Determinants of patterns of regional morphotype pools

We identified 23 morphotypes (12 detachable leaf, 9 stem, and 2 integral leaf). Cynipid morphotype community composition varied among regions (Fig. 2). Variation in CA1 is influenced by detachable leaf gall and stem gall types that dominate in R1-3 and are lacking in the expanded range, which is dominated by integral leaf gall types (R5-6) (Appendix S1: Table S5). Detachable cynipids have negative loadings along PC1 (*Trichoteras tubifaciens* = −0.97, *Andricus stellaris* = −0.92, *A. kingi* = −0.85), as do stem cynipids (*A. chrysolepidicola* = −0.30, *A. coortus* = −0.27, *A. quercuscalifornicus* = −0.27) while integral leaf gall formers (*N. washingtonensis* = 0.26, *A. opertus* = 0.40) have positive loadings*. N. saltatorius* has a positive loading along CA1 (0.66), reflecting its high abundance in the expanded range. Two other detachable cynipids were found in the expanded range - *Besbicus mirabilis* is common and *A. kingi* found at one site. CA2 variation is influenced by rare species (“pink bowtie” = −0.65 and *Disholcaspis canescens* = 0.93).

Latitude was retained in all the final models (Fig. 3 A-D, Table 1; Appendix:Table S8). Cynipid richness followed the predicted negative relationship with increasing latitude (*F* = 72.66, *P* < 0.0001; Fig. 2A). CA1 increased with latitude (*F* = 126.27, *P* < 0.0001, Fig. 3B), revealing latitudinal changes in community composition. Gall abundance excluding *N. saltatorius* was negatively related to latitude (*F* = 6.47, *P* = 0.065, Fig. 2C). However, *N. saltatorius* abundance and total gall abundance had a positive relationship with latitude (*N. saltatorius* abundance: *F* = 19.48, *P* <0.0001, Fig. 2D; total gall abundance: *F* = 3.89, *P* = 0.054, Fig. S5) (Table 1). The highest abundances of *N. saltatorius* were in R5 with approximately 20,000 more galls compared to other regions (Appendix S1:Fig. S6). PC1temp was retained in several final models (Table 1) and has a strong positive correlation with latitude (*R* = 0.92) such that higher latitude sites had lower maximum and minimum temperatures and shorter seasonality (Appendix S1:Table S3). Elevation was negatively related to latitude (*R* = −0.71) and had a negative relationship with richness (Table 1). Total oak area and PC2_habitat_ have a negative relationship with richness with oak area negatively related to latitude (*R* = −0.78) and PC2_habitat_ positively related (*R* = 0.77) (Appendix S1:Table S3).

**Figure 3:**
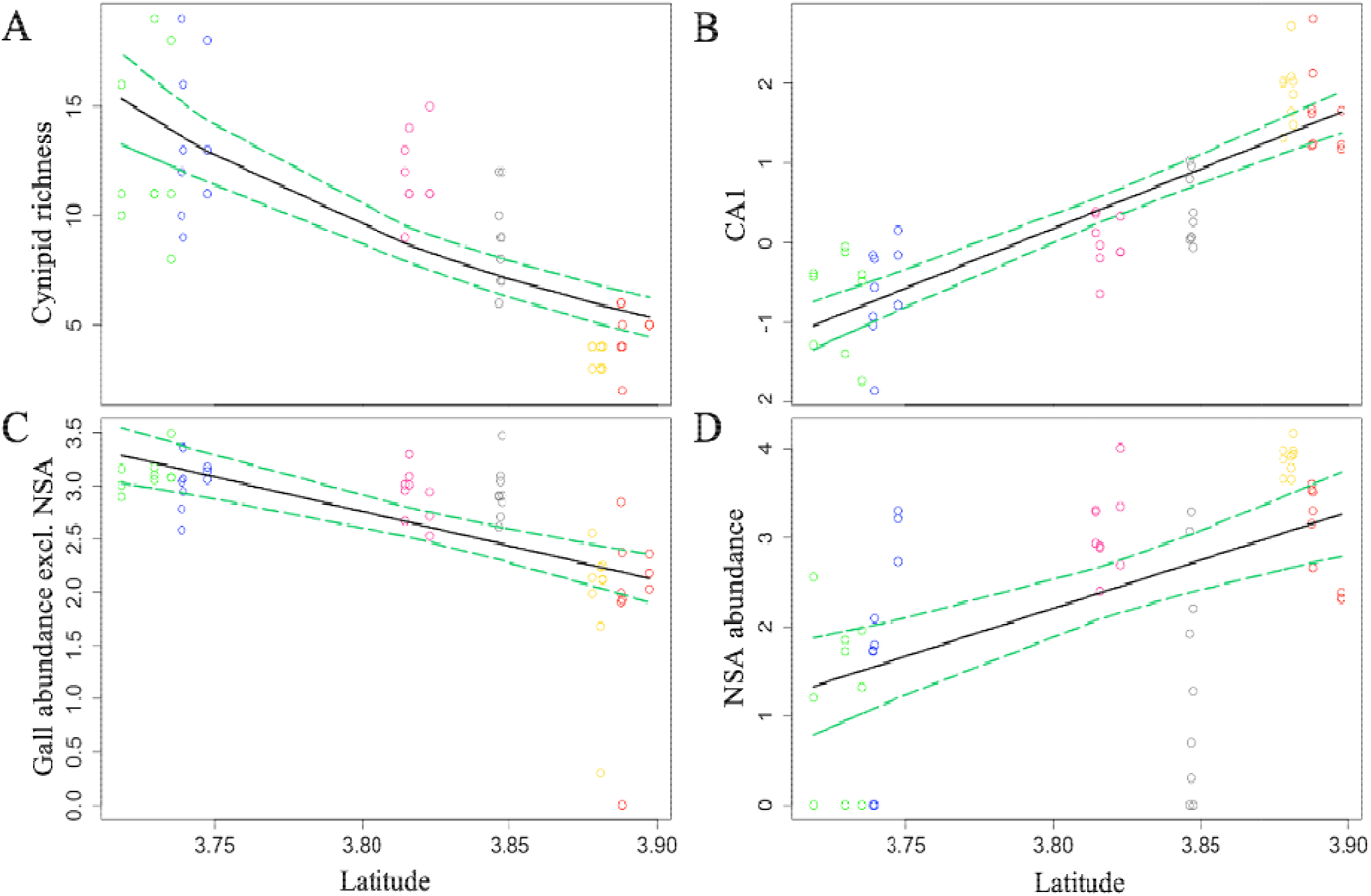
Regressions between a) cynipid morphotype richness, b) cynipid morphotype composition (CA1), c) gall abundance excluding *N. saltatorius* (log-transformed), and d) *N. saltatorius* abundance (log-transformed) and latitude (log-transformed) for each survey date/site. Regions are colored coded (Fig. 1). Solid black line represents predicted regression lines and 95% C.I (green dashed lines) for fixed effects (latitude) only for clarity (Table 1).

**Table 1:** Slopes for best fit models after Akaike information criterion (AICc) comparisons of the final model sets. Within parentheses are confidence intervals for slopes. Bolded slopes are significant. See AICc output in Appendix 1:Tables S6-8.

| Predictor variables | Cynipid morphotype richness | Cynipid community Composition (CA1) | Total gall abundance | Gall abundance excluding *N. saltatorius* | *N. saltatorius* abundance |
| --- | --- | --- | --- | --- | --- |
| Latitude | **-5.58 (± 2.00)** | **14.55 (± 2.78)** | **-6.46 (±1.74)** | -5.79 (±2.28) | -1.99 (± 5.75) |
| Elevation | **-0.64 (± 0.18)** |  |  |  |  |
| PC1<sub>temp.</sub> | **-0.27 (± 0.11)** |  | **0.36 (± 0.07)** |  | **0.55 (± 0.23)** |
| PC2<sub>habitat</sub> | -0.19 (±0.13) | | | | |
| Oak habitat area | **-0.0003 (± 0.18)** |  |  |  |  |

### Patterns in local morphotype interactions

The number of significant cynipid morphotype associations varied among regions (Fig. 4). Most significant associations occurred between the diverse detachable leaf cynipids (including *N. saltatorius*), while stem and integral cynipids had fewer significant associations. Across regions, stem cynipids had 8, integral leaf 5, and detachable leaf 58 significant associations. There was a decrease in significant associations with increasing latitude. *N. saltatorius* associates with 4 species directly and 3 indirectly in R1, 4 directly and 4 indirectly in R2, 1 directly, and 0 indirectly in R3, and 2 directly and 2 directly in R4, but had no significant associations in R5 and R6. All direct associations of *N. saltatorius* are with detachable leaf cynipids in the native range (Fig. 2). Local cynipid patterns were influenced by multiple factors, mainly Julian date and soil moisture in R1-R4 but not for R5-R6 (Appendix S1:Fig. S7, Table S9).

**Figure 4:**
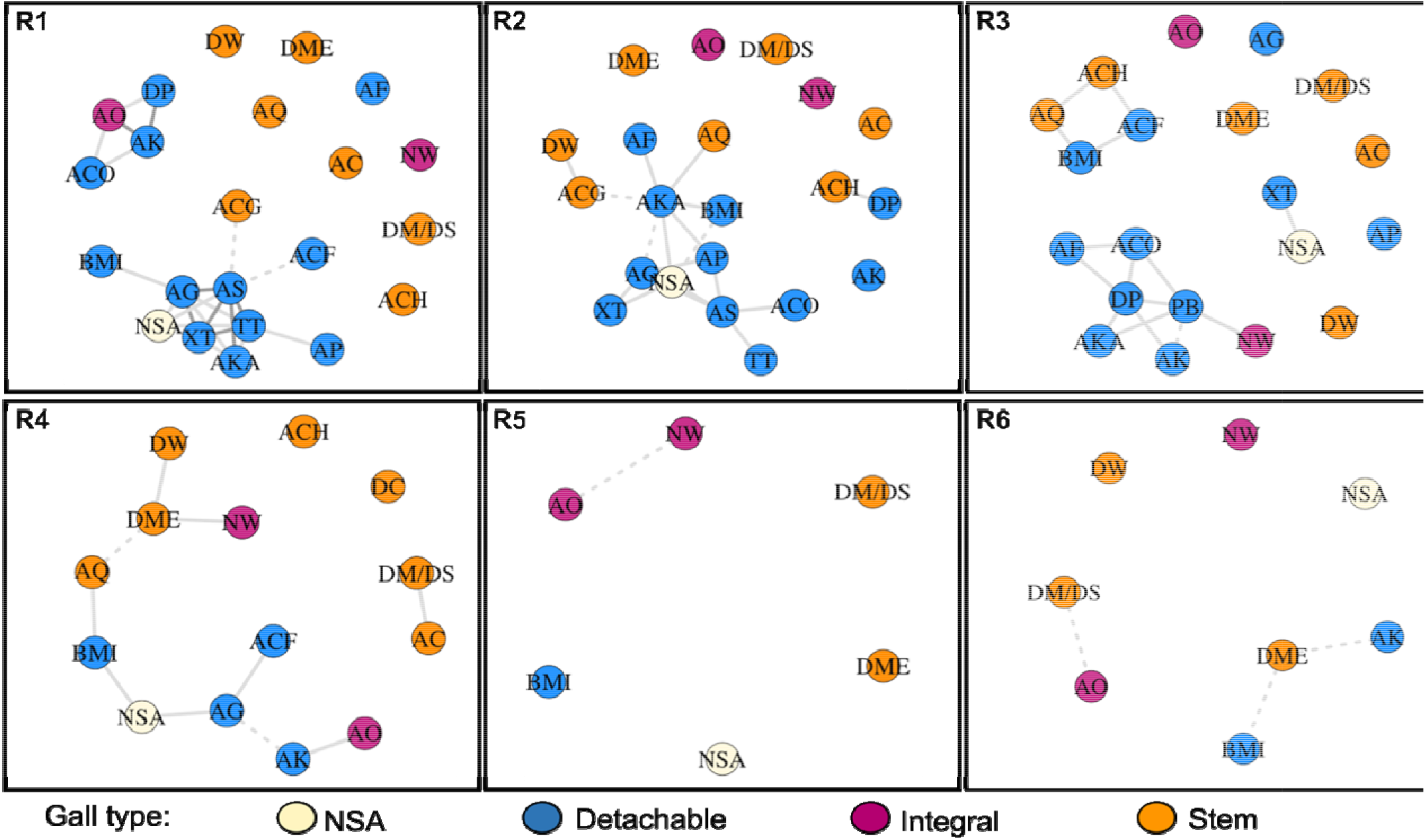
Partial correlation analysis using presence absence data of cynipid morphotypes on trees. Networks display interactions between morphotypes on trees (n = 90) in a region. Shades of lines represent correlation strength (weak (0.25-0.49), moderate (0.5-0.74), and strong (0.75-1.0)). Solid lines indicate a significant correlation while dashed lines are non-significant (Spearman test). Morphotypes colored by gall type category (blue = detachable, purple = integral, orange = stem), morphotype abbreviations found in Table S2.

*N. saltatorius* gall abundance had a significant negative relationship with detachable leaf gall abundance (*F* = 4.19, *P* = 0.04; Fig. 5a) along with integral leaf gall abundance (*F* = 15.92, *P* < 0.01; Fig. 5b). *N. saltatorius* abundance had no relationship with stem gall abundance (*F* = 0.58, *P* = 0.446; Fig. 5c). For leaf and integral gall abundance there was also a significant effect of region (leaf: *P* < 0.0001, integral: *P* < 0.0001); but no significant interactions between region and gall type abundance for any of the gall types (leaf: *P* = 0.112, integral: *P* = 0.604).

**Figure 5:**
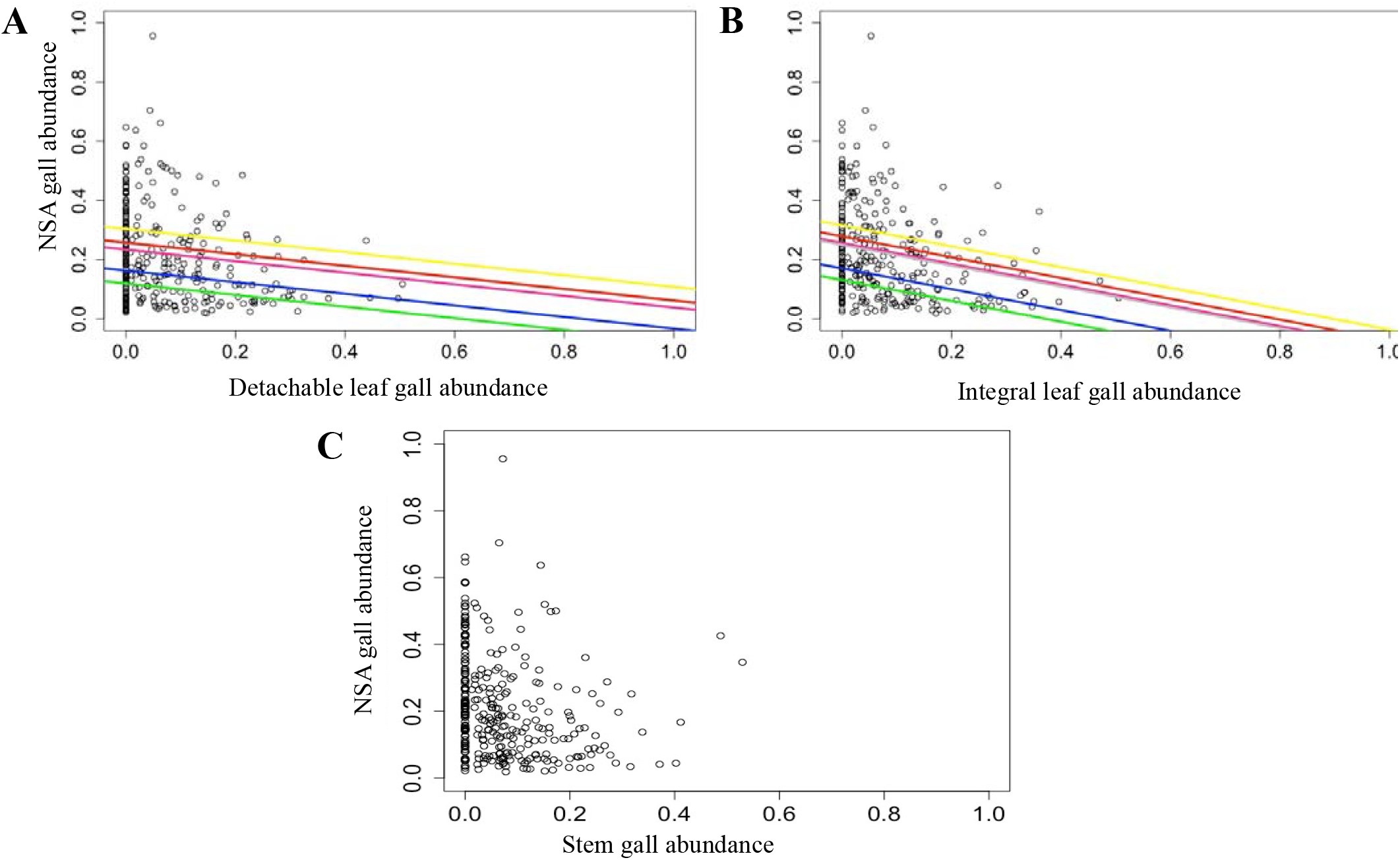
Regressions between relative abundance of a) other detachable leaf cynipids, b) integral leaf cynipids, and c) stem cynipids and *N. saltatorius* abundance. Symbols represent trees where *N. saltatorius* and gall types co-occur. Gall abundances are transformed (arcsine sqrt). Regression lines (LM) are shown for significant relationships. Colors represent regions (Fig. 1).

## Discussion

As predicted, cynipid diversity followed an LDG, with fewer morphotypes at higher latitude sites. Community composition changed with a loss of detachable leaf and stem gall types at high latitudes. These regional diversity patterns are correlated with gradients in abiotic factors, such as variation in temperature extremes, and habitat factors, such as oak patch size, that correlate with latitude. At local scales (on trees) *N. saltatorius* interacted the most with other detachable leaf cynipids that were largely absent in the expanded range, where only 2 (out of 12) morphotypes were present. On trees where other detachable leaf cynipids and *N. saltatorius* co-occurred, their abundances were negatively related, suggesting antagonistic interactions. Our results indicate that LDGs of oak cynipid communities provide high niche opportunities in northern oak populations, resulting from weak local antagonistic interactions.

Phytophagous insect communities, including cynipids, that occupy post-glacial landscapes in the northern hemisphere established through migration after glaciers retreated, with source refuges often coming from lower latitudes (Hayward and Stone 2005, Mutun 2016, Stone et al. 2017). *Q. garryana* maintained a large portion of its current range during the last glacial maximum (LGM) (Fraser glaciation period, 15 kyr BP), including northern populations in Oregon and Washington (Booth et al. 2003). The only portion of the current range of *Q. garryana* that was glaciated during the LGM was the Puget Sound region, San Juan Islands, WA and Vancouver Island BC (Brubaker 1991, Brown and Hebda 2002). *Q. garryana has* reduced genetic diversity in northern edges and on islands, suggesting founder effects as it expanded its range after ice sheets receded (Marsico et al. 2009). As *Q. garryana* experienced bottlenecks when expanding, so may have their associated insect communities. In other regions, there are latitudinal trends in cynipid communities with expansion from refuges occurring post-glaciation (Hayward and Stone 2005, Mutun 2016), with greater community (and genetic) diversity in the core, reducing towards the poles (Stone et al. 2017). The process of cynipid dispersal over long distances is not fully understood as they are weak fliers and dispersal limited, but wind is a possible mechanism, along with accidental transport to neighboring ranges or over long distances (Gilioli et al. 2013).

Post-glaciation dispersal lags and limitations likely partially determine current cynipid distributions. In this case, lower diversity in northern latitudes is confounded with an island effect; however, we see lower diversity and changes in composition in R3 and R4, compared to R1 and R2 suggesting a latitudinal gradient on the mainland. Host plants and ecosystems are often patchier at range edges, which is the case for *Q. garryana-ecosystems* both on the mainland in Washington and the Island (Fuchs 2001, Loughnan and Williams 2019). Dispersal limitation across unsuitable non-oak habitat (forests) or other barriers, such as topographical or water barriers could contribute to LDGs (Gaston 2000, Willig et al. 2003, Kerr et al. 2016). Patch size correlated negatively with latitude and cynipid richness, where conditions are cooler with higher precipitation - more conducive to *P. menziesii* forests. In addition to limited dispersal to more isolated patches, small patches could contribute to reduced patterns in richness, as larger patches often harbor higher species richness (Steffan-Dewenter and Schiele 2008). Patches are also fractured due to land use change, especially in BC as most suitable habitat is encroached by urban and suburban development (Fuchs 2001, Loughnan and Williams 2019). However, we did not find patterns in habitat and land use borders (other than the percent of oak border) contributed to patterns in richness and abundance.

The other regional factor that correlated with latitude and explained patterns in cynipid richness was PC1_temp_, representing temperature extremes and seasonality. Studies have shown that gall formers are more common in dry and warm environments (Gonçalves-alvim and Fernandes 2001), which represent the conditions of southern populations of *Q. garryana*.Minimum temperatures decrease with increasing latitude and elevation, and studies have proposed that climatic factors may limit phytophagous insects’ distribution and diversity at high latitudes and elevations (Willig et al. 2003, Wieski and Pennings 2014, Fitt and Lancaster 2017, Zvereva and Kozlov 2021). Since PC1_temp_ correlated highly with latitude, we do not know the extent to which temperature contributes to patterns in diversity.

We found high abundances of *N. saltatorius* at northern latitudes, in the expanded range, with no other factors contributing to this pattern. Thus, landscape and abiotic factors that limit cynipid richness and abundance at high latitudes, do not seem to apply to the range-expanding species. Thus, patterns in *N. saltatorius* abundances may be driven by other factors, such as release from biotic interactions (Prior and Hellmann 2013, Fitt and Lancaster 2017).

*N. saltatorius* underwent a poleward range expansion, but it is unclear whether this is linked to climate change or an introduction. Regardless, poleward range expansions occur via various types of anthropogenic change, where interacting species are not expected to move in concert (Gilman et al. 2010, Wallingford et al. 2020). One difference between populations undergoing climate-driven versus other types of range expansions is that in the former case, expanding populations may come from populations at the edge of their thermal tolerances. To this end, populations moving into the expanded range be small and limited by abiotic conditions. Alternatively, if edge (source) populations are locally adapted to edge conditions, their populations might be larger and less limited by expanded range (new edge) conditions (Hellmann et al. 2008, Pelini et al. 2009). The high abundance of *N. saltatorius* in its expanded range suggests that abiotic conditions in the expanded region are not limiting.

Cynipids are patchily distributed within oak populations, with some trees lacking galls while others have moderate or high abundances (Egan and Ott 2007, Prior and Hellmann 2010). We found that detachable leaf cynipids had the strongest associations on trees, including with our focal species, while other gall types largely lack strong associations. Since only two detachable leaf cynipids occur in R5 and R6, *N. saltatorius* does not have strong associations on trees in its expanded range. Several factors could contribute to gall associations (or similar preferences) for individual trees. Julian date contributed to variation in association among trees (R1-4), which is expected given our sampling over time. Soil moisture and temperature (that likely correlate with host plant phenology) explained trends, suggesting that microhabitat differences influence gall associations. There are factors that were beyond the scope of this study to measure that likely contributes to cynipid associations, such as host plant genetic variation and variation in defenses.

Sedentary insects interact on their host plants (Veen et al. 2006, Kaplan and Denno 2007, Prior and Hellmann 2013, Holt and Bonsall 2017). Our predictions that weak associations might lead to high niche opportunities assumes that weak associations on trees release *N. saltatorius* from antagonistic interactions. However, cynipids may coexist on trees while not antagonistically interacting. The negative relationship between *N. saltatorius* abundance on trees and the abundance of detachable and integral gall wasps, but not with stem gall wasps, suggests antagonistic interactions. Future studies might manipulate competitor presence of abundance to examine how this affects *N. saltatorius* survival and measure the proportion of shared enemies between gall morphotypes.

That *N. saltatorius* mostly associates with detachable leaf cynipids on trees, but negatively interacts when co-occurring, suggests that release from smaller regional pools that lack the detachable gall group might contribute to ecological release. Cynipids can compete indirectly via altering resources by acting as nutrient sinks or sequester carbon, or they could interact through induced or constitutive defenses (Cuevas-Reyes et al. 2003, Prior and Hellmann 2010). Mechanisms of host defenses against cynipids are not well understood. Chemical defenses against chewing herbivores in oaks (tannins, lignin) is suspected to not be a main defense against galling insects (Barbosa and Fernandes 2014). A hypersensitive response, in which tissue surrounding galls necrose or whole leaves drop, has been described as a defense against gall formers (Barbosa and Fernandes 2014). If induced, this response could impede other individuals from establishing, but little is known about mechanisms of this defense mechanism against galling insects. While we found negative correlations in abundances between species suggesting antagonistic interactions, species could also compete over ‘evolutionary time’ such that lower diversity at the poles might have created relaxed selection for defensive responses particularly against detachable leaf galls in northern populations (Woods et al. 2012, Moreira et al. 2015).

Insects also compete through shared enemies, i.e., apparent competition. We expect stronger apparent competition for insect hosts that have similar morphological and phenological traits (Bailey et al. 2009, Schönrogge et al. 2012, Holt and Bonsall 2017). *N. saltatorius* not only shares the locality at which it forms its galls with other leaf detachable gall wasps, but it also shares similar morphological and phenological traits (Prior, unpublished data). For example, detachable leaf galls overlapped more in time with *N. saltatorius*, and many of these galls are small, single chambered galls lacking gall defenses such as hard woody exteriors and large interstitial space. Galls with similar shapes and phenology are more likely to share parasitoids (Bailey et al. 2009). *B. mirabilis*, the predominant detachable leaf gall on the Island, does not share many parasitoids with *N. saltatorius* (Prior, unpublished data), and it morphologically different (25X larger). It is important to note that the spatial scale for apparent competition might be greater than species co-occurring on trees, since parasitoids are not sessile (Holt and Bonsall 2017). However, surveyed trees were spread throughout sites, and we assume co-occurring species are most likely to experience apparent competition.

Leveraging anthropogenic range expansions, is a powerful approach to uncover the community dynamics of range expansions. This research provides valuable insight into how biogeographical patterns in communities and species interactions can influence the outcome of range expansions. By studying cynipid communities across the entire range of *Q. garryana*, we uncovered that diversity of cynipid communities across their range creates high niche opportunities (or weak biotic interactions) that might contribute to ecological release. Uncovering how regional and local scale patterns and processes combine to contribute to niche opportunities is important for uncovering the role of biotic interactions in range expansions. Given the ubiquity of LDGs, these outcomes might be common in other phytophagous insect communities.

## Acknowledgements

We thank Katie Harms for help in the field, Catherine Ruis, Leslie Huang, Jesse Lofaso, Kelly McGourty, Julia Berliner, Sage Daughton, and Serena Feldman for help in the lab. We thank landowners: CA BLM, OR Parks & Rec. Dept., OR Dept. of Fish & Wildlife, US Forest Service, The Nature Conservancy, City of West Linn Park, Southern OR Land Conservancy (private owner Stan Dean), WA DNR, Weyerhaeuser, Canada Dept. National Defense, Saanich Parks, BC Parks, Nature Conservancy Canada. Funding was provided by National Geographic Society (NGS-53395R-18) (KMP), National Science Foundation (DEB 1934387) (KMP), Clark Fellowship (DGJ), and Binghamton University. Research was conducted on stolen land belonging to Indigenous tribes and people that historically and present day occupy lands: Á,LE□ENE□ □TE (W□SÁNEĆ), Quw’utsun, Ts’uubaa-asatx, Stz’uminus, sc□әwaθena□ □ tәmәx□ (Tsawwassen), Atfalati, Chehalis, Coast Salish, Cow Creek Umpqua, Cowlitz, K’ómoks, Kalapuya, Lekwungen/Songhees, Multnomah, Nisqually, S’Klallam, Sasha, Snuneymuxw, Snaw-naw-as, Modoc, Takelma, Tolowa Dee-ni’, Wasco and Wishram, Yakama, Karuk.

## Author contributions

KMP, DGJ, THQP developed ideas, DGJ, JK, KMP implemented field protocols, DGJ analyzed data with input from KMP, JMR performed habitat analysis, DGJ, KMP, JK wrote manuscript with input from all authors.

## Data and code available

https://doi.org/10.6084/m9.figshare.18737834.v1

## Notes

https://doi.org/10.6084/m9.figshare.18737834.v1

### Competing Interest Statement

The authors have declared no competing interest.

