## Supplemental Methods, Tables, & Figures for "Latitudinal gradient in species diversity provides high niche opportunities for a range-expanding phytophagous insect"

**Appendix S1**

**Latitudinal gradient in species diversity provides high niche opportunities for a range-expanding phytophagous insect**

Authors: Dylan G. Jones; Julia Kobelt; Jenna M. Ross; Thomas H.Q. Powell; Kirsten M. Prior

**Section S1: Supplementary Methods**

*Measuring abiotic and habitat variables*

We hung an iButton temperature logger (1-Wire, Thermochron) on a reachable branch (approximately 1 m and in the canopy, out of direct sunlight) on a survey tree not located at the edge of a site on the first survey day. Loggers were kept up until the final sampling date for each site. For each survey tree, we measured tree height (m) and diameter at breast height (DBH), soil temperature and soil moisture at the base of the tree. By placing a transect tape at the base of the tree and walking along the flattest surface until we could see the top of the tree. We then determined the angle from where we were standing to the top of the tree, and calculated tree height (m). Diameter at breast height (DBH) was measured using a girthing tape (cm). Soil temperature (°C) and moisture (qualitative moisture scale 1-10; 1-3: dry, 4-7: moist, 7-10: wet) were measured by taking the average of three locations around the base of each tree using temperature and soil moisture probes. Since tree height and DBH were correlated (*R* = 0.71), we created a composite ‘tree size’ variable as PC1 from a principal component analysis (PCA).

For each site, we collected BioClim variables with a spatial resolution of ~340 km^2^ from WorldClim 1970-2000 data: temperature (annual mean temperature, temperature maximum, temperature minimum, and temperature seasonality), and precipitation (annual precipitation, precipitation minimum, precipitation maximum, precipitation seasonality) (Fick 2017) and created composite climate variables using PCA representing temperature and precipitation separately (Gonçalves-Souza et al. 2014, Moreira et al. 2014, 2015, Romero et al. 2016). We chose these variables as they are often contributors to patterns in insect species distributions along latitudinal gradients (Gonçalves-Souza et al. 2014; Moreira et al. 2014, 2015; Romero et al. 2016, Lembrechts et al. 2019). PC1_temp_ reflected variation in extremes and seasonality (max. temp., min. temp., and temp. seasonality), and PC2_temp_ reflected variation in annual mean temperature (Fig. S1a). PC1_precip._ reflected min., max., and annual precipitation, while PC2_precip_ reflected precipitation seasonality (Fig. S1b). PC1_temp_ was strongly correlated with latitude (*R* = 0.92), followed by PC1_precip_ (*R* = 0.66). PC2_temp_ (*R* = 0.17) and PC2_precip_ (*R* = 0.29) were not correlated with latitude (Table S3).

We loaded coordinates of survey trees into Google Earth to calculate the sampling area (m^2^) encompassed by the surveyed trees at each site and to define habitat and land use surrounding sites by creating a 1 km extended permitter to classify broad categories. We measured a 1-kilometer extended perimeter around survey areas and used satellite images to classify the extended perimeter into broad visible habitat and land use categories: oak savanna/oak woodland (dominated by oaks), forest (high density canopy cover areas, often dominated by *P.* *menziesii*), urban or suburban land use (*hereafter* urban), grassland or old field (*hereafter* grassland), active agricultural field, water body, and island along with the percent area of each habitat and land use types. We performed a PCA to create composite variables representing survey perimeter habitat. PC1_habitat_ represented variation in non-oak habitat and land use types, with sites associated with field, urban, and forest habitats associated with negative values and with island, water, field, and urban associated with positive values (Fig S2). To estimate oak patch size, we summed the oak survey area and the oak habitat perimeter area. This underestimates oak habitat which is continuous in the south and patchy in the north, but captures that oak patches are larger in southern locations. We also estimated distance to edge for each tree as the distance of each surveyed oak tree to the nearest non-oak habitat or land use type. We extended a radius from each tree until it reached the closest non-oak habitat or land use type or extended to 1,000 meters. PC2_habitat_ was positively correlated with latitude (*R* = 0.77) and total oak area negatively correlated with latitude (*R* = -0.78). Survey area and PC1_habitat_ were weakly correlated with latitude (Table S3).

*Regional analysis and mixed effect models*

First, we ran single mixed effect models where each dependent variable was paired with a single predictor variable and calculate Akaikie information criterion (AICc) (Vasconcelos et al. 2018). We then built models by creating multi-factor models using a forward step-wise approach starting with variables with the two lowest AICc scores, and then building to add in all variables (we did not include interactions). First, we created and compared models with geographic + abiotic variables (14 models, including all single and multi-factor models) (Table S6). Then we created and compared models with geographic + habitat variables (14 models) (Table S7). We used the AICcmodavg package in R to compare AICc scores among models in sets (Mazerolle 2016). We retained all models for each set that resulted in a ΔAICc < 2 (Hill et al. 2017; Cruaud et al. 2018; Loughnan and Williams, 2019). Then, we created a final model set for each response variable including retained predictor variables (either in a single or a multi-model), building to full models in a forward stepwise approach as above (Table S8). We performed model comparisons for our final model set as stated above. We used the R package LmerTest to obtain slope estimates and p-values for models with a ΔAICc < 2 (Kuznetsova et al. 2017). For abundances (total cynipid abundance, abundance of *N. saltatorius*, and abundance of all cynipids – without *N. saltatorius*) and CA1 and CA2 (see main text) we performed LMM, performing log transformations on gall abundances. For richness, we ran a GLMM (negative-binomial). We used the package lme4 in R to perform generalized linear mixed-effects models (Bates et al. 2020).

*Factors influencing local cynipid composition* (constrained canonical analysis, CCA)

To determine what abiotic or habitat factors influence cynipid composition on trees, we performed a constrained canonical analysis (CCA) with presence/absence data on each tree pooled for each region (Oksanen et al. 2013). We used the following predictor variables that we measured within sites: Julian date, corrected soil temperature, corrected soil moisture, distance to closest non-oak border, and tree size. Because soil temperature is affected by survey time and date, we perform a regression between soil temperature and ambient temperature (from Ibuttons at the site) at the same date and time point the tree was surveyed. We used the residuals for each tree replicate as corrected temperature. We also performed a similar linear regression to correct for soil moisture using Julian date.

Veech, Joseph A. and Crist, Thomas O., PARTITION: software for hierarchical partitioning of species diversity, version 3.0, 2009.

**Section S2: Supplementary Figures and Tables**

**Table S1**: Study site information along with survey dates, size of survey area, and predominant surrounding broad habitat category type.

| **Region** | **Site acronym** | **Site Name/Owner or Manager** | **Latitude**  **/Longitude** | **Survey 1** | **Survey 2** | **Survey 3** | **Survey area (m^2^)** | **Dominant surrounding habitat** |
| --- | --- | --- | --- | --- | --- | --- | --- | --- |
| 1 | ASH | **Ash Creek Road**  Bureau of Land Management | 41.8891,  -122.5820 | 5/24/19 | 6/13/19 | 7/6/19 | 73874 | Oak |
| 1 | COP | **Copoco Lake**  Bureau of Land Management | 41.2054,  -123.9517 | 5/21/19 | 6/6/19 | 7/4/19 | 54824 | Oak |
| 1 | ICR | **Indian Creek Road**  Bureau of Land Management | 41.6583,  -122.8529 | 5/23/19 | 6/12/19 | 7/5/19 | 102975 | Oak |
| 2 | COL | **Colestin Valley**  Southern Oregon Land Conservancy, private landowner | 42.0447,  -122.6431 | 6/2/19 | 6/25/19 | 7/10/19 | 60007 | Forest |
| 2 | STAR | **Star Ranger**  US Forest Service | 42.0665,  -123.1104 | 5/26/19 | 6/17/19 | 7/8/19 | 70681 | Forest |
| 2 | WHET | **Whetstone Preserve**  The Nature Conservancy | 42.4155,  -122.9097 | 5/27/19 | 6/18/19 | 7/9/19 | 118772 | Agriculture |
| 3 | LINN | **West Linn Oak Savanna**  The Trust for Public Land, City of West Linn Parks and Recreation | 45.3511,  -122.6517 | 5/30/19 | 6/22/19 | 7/16/19 | 45828 | Urban |
| 3 | ME | **Memaloose State Park**  Oregon Parks and Recreation Department | 45.4145,  -121.2017 | 6/8/19 | 6/23/19 | 7/17/19 | 188671 | Oak |
| 3 | OI | **Oak Island**  **Sauvie Island Wildlife Area**  Oregon Department of Fish & Wildlife | 45.7248,  -122.8181 | 5/29/19 | 6/21/19 | 7/15/19 | 928419 | Water |
| 4 | BAH | **Bald Hills, Vail Tree Farm**  Weyerhauser | 46.8090,  -122.4356 | 6/5/19 | 6/30/19 | 7/2019 | 13763 | Forest |
| 4 | GHE | **Glacial Heritage Preserve**  Thurston County Parks; Washington Department of Fish and Wildlife | 46.8632  -123.0379 | 6/6/19 | 7/1/19 | 7/19/19 | 53591 | Forest |
| 4 | SC | **Scatter Creek Wildlife Area**  Washington Department of Fish and Wildlife | 46.8332,  -122.9991 | 6/7/19 | 6/29/19 | 7/21/19 | 53591 | Field |
| 5 | LMD | **Mount Douglas Park**  **(Little Mount Douglas)**  Saanich Parks | 48.4907,  -123.3514 | 6/16/19 | 6/27/19 | 7/23/19 | 31969 | Agriculture/  Urban |
| 5 | MTO | **Mount Tolmie Park**  Saanich Parks | 48.4583,  -123.3224 | 6/15/19 | 6/26/19 | 7/23/19 | 40049 | Urban |
| 5 | RP | **Rocky Point**  Department of National Defense | 48.3230,  -123.541 | 6/17/19 | 7/4/19 | 7/26/19 | 201197 | Water  /Forest |
| 6 | COW | **Cowichan Nature Preserve**  Nature Conservancy Canada | 48.8075,  -123.631 | 6/19/19 | 7/3/19 | 7/24/19 | 46444 | Agriculture  /Urban |
| 6 | MTZ | **Mount Tzouhalem Ecological Reserve**  British Columbia Parks | 48.7890,  -123.6368 | 6/24/19 | 7/8/19 | 7/25/19 | 22999 | Forest |
| 6 | NAN | **Notch Hill**  Department of National Defense | 49.2715,  -124.1561 | 6/20/19 | 7/9/19 | 7/29/19 | 20573 | Forest |

**Table S2**: List of oak cynipid morphotypes identified during survey periods, morphotype acronyms, gall type group, if galls are known to be polythalamous (contain multiple chambers) or monothalamous (one chambers) and regions and sites with where morphotypes were collected (see Table S1 for site acronyms).

| **Scientific Name**  **Common Name** | **Acronym** | **Gall type** | **Regions** | **Region (Sites)** |
| --- | --- | --- | --- | --- |
| ***Andricus coortus***  Club gall wasp | *AC* | Stem (petiole),  Polythalamous | 1-4,5 | 1 (ASH, COP, ICR); 2 (COL, STAR, WHET) 3(LINN, ME, OI), 4 (BAH, GHE, SC) |
| ***Andricus confertus***  Convoluted gall wasp | *ACF* | Leaf (detachable),  Monothalamous | 1,3,4 | 1 (ASH); 3 (LINN, OI); 4 (BAH, GHE, SC) |
| **Unknown species**  Acorn cap gall | *ACG* | Stem (acorn) | 1,2 | 1 (ICR); 2 (WHET) |
| ***Andricus chrysolepidicola***  Irregular spindle gall wasp  (agamic) | *ACH* | Stem (integral),  Polythalamous | 1-4 | 1 (ASH, COP); 2 (COL, STAR, WHET); 3 (LINN, OI) 4 (BAH, GHE, SC) |
| Unknown species  Acorn gall | *ACO* | Stem (acorn) | 1-3 | 1 (ASH, COP, ICR); 2 (COL); 3 (LINN, ME) |
| ***Andricus fullawayi***  Yellow wig gall wasp | *AF* | Leaf (detachable),  Monothalamous | 1-3 | 1 (ICR); 2 (STAR, WHET); 3 (ME) |
| ***Andricus gigas***  Saucer gall wasp  (agamic) | *AG* | Leaf (detachable),  Monothalamous | 1-4 | 1 (ASH, COP, ICR); 2 (COL, STAR, WHET); 3 (ME, OI); 4 (BAH, GHE, SC) |
| ***Andricus kingi***  Red cone gall wasp  (gamic, agamic) | *AK/*  *AKA* | Leaf (detachable),  Monothalamous | 1-3,4,6 | 1 (ASH, COP, ICR); 2 (COL, STAR, WHET) 3 (LINN, ME, OI); 4 (BAH, GHE, SC); 6 (MTZ) |
| ***Andricus opertus***  Fimbriate gall wasp | *AO* | Leaf (integral),  Monothalamous | 1-6 | All sites |
| ***Andricus parmula***  Disc gall wasp | *AP* | Leaf (detachable),  Monothalamous | 1-3 | 1 (ICR); 2 (COL, STAR, WHET); 3 (LINN, ME, OI) |
| ***Andricus quercuscalifornicus***  California gall wasp | *AQ* | Stem (integral),  Polythalamous | 1-4 | 1 (ASH, COP, ICR); 2 (COL, STAR, WHET); 3 (LINN, ME, OI); 4 (BAH, GHE, SC) |
| ***Andricus stellaris***  Stellar gall wasp | *AS* | Leaf (detachable),  Monothalamous | 1,2 | 1 (ASH, COP, ICR); 2 (COL, STAR) |
| ***Besbicus mirabilis***  Speckled gall wasp | *BMI* | Leaf (detachable).  Monothalamous | 1-6 | 1 (ASH, COP, ICR); 2 (COL, STAR, WHET); 3 (LINN, ME, OI); 4 (BAH, GHE, SC) 5 (LMD, MTO, RP); 6 (COW, MTZ) |
| ***Disholcaspis canescens***  Round honeydew gall wasp | *DC* | Stem (detachable),  Monothalamous | 4,6 | 4 (BAH); 6 (MTZ, NAN) |
| ***Disholcaspis mamillana/***  ***Disholcaspis simulata***  Bullet gall wasp  /Peach gall wasp | *DM/DS* | Stem (detachable),  Monothalamous | 1-6 | 1 (ASH, COP, ICR); 2 (COL, STAR, WHET); 3(LINN, ME, OI); 4 (BAH, SC); 5 (MTO, MTZ, NAN) |
| ***Disholcaspis mellifica***  Flat-topped honeydew  /Twig gall wasp | *DME* | Stem (detachable),  Monothalamous | 1-6 | 1 (ASH, COP, ICR); 2 (COL, STAR, WHET); 3 (LINN, ME, OI); 4 (BAH, GHE, SC); 5 (LMD, MTO, RP); 6 (COW, MTZ, NAN) |
| ***Andricus (Dros) pedicellatum***  Hair streak gall wasp | *DP* | Leaf (detachable),  Monothalamous | 1-4 | 1 (ASH, COP, ICR); 2 (COL, STAR, WHET); 3 (ME, O1) |
| ***Burnettweldia (Disholcaspis) washingtonensis***  Fuzzy gall wasp | *DW* | Stem (detachable),  Monothalamous | 1-3,4,6 | 1 (COP, ICR, COL); 2 (WHET); 3 (LINN, OI) 4 (BAH, GHE, SC) 6 (MTZ, NAN) |
| ***Neuroterus saltatorius*** Jumping all wasp | *NSA* | Leaf (detachable),  Monothalamous | 1-6 | All sites |
| ***Neuroterus washingtonensis***  Midrib gall wasp | *NW* | Leaf (integral),  Polythalamous | 1-6 | All sites |
| **Undescribed**  Pink bowtie gall wasp | *PB* | Leaf (detachable),  Monothalamous | 3 | 3 (ME) |
| ***Andricus (Trichoteras) tubifaciens***  Crystalline tube gall wasp | *TT* | Leaf (detachable),  Monothalamous | 1,2 | 1 (ASH, COP, ICR) 2 (STAR) |
| ***Xanthoteras teres***  Ball-tipped gall wasp | *XT* | Leaf (detachable),  Monothalamous | 1-3 | 1 (ASH, COP, ICR); 2 (COL, STAR, WHET); 3 (LINN, ME, OI) |

**Table S3**: Predictor variables used in models to measure factors influencing latitudinal patterns in regional diversity. *R* values represent correlations of each variable with latitude.

| **Predictor variable** | **Correlation with latitude (*R* value, direction)** | **Variable Description** |
| --- | --- | --- |
| **Geographical variables** | | |
| **Latitude** | n/a | Latitude of study site (log-transformed) |
| **Elevation** | 0.71 (-) | Elevation of study site (m) (log-transformed) |
| **Abiotic variables** | | |
| **PC1_temp_** | 0.91(-) | PC1 of temperature BIOCLIM variables. Positive values represent lower temperature seasonality, low maximum temperatures, and high minimum temperature at sites (Fig S1a). |
| **PC2_temp_** | 0.17 (-) | PC2 of temperature BIOCLIM variables. Positive values represent greater annual mean temperature at sites (Fig. S1a). |
| **PC1_precip._** | 0.66 (+) | PC1 of precipitation BIOCLIM variables. Positive values represent greater maximum precipitation, lowest precipitation, annual precipitation at sites (Fig. S1b). |
| **PC2_precip_** | 0.29 (+) | PC2 of precipitation BIOCLIM variables represent lower seasonality at sites (Fig S1b). |
| **Soil moisture** | 0.19 (-) | Residuals between average soil moisture among trees at a site during a survey period and average air temperature on the survey day. |
| **Habitat variables** | | |
| **PC1_habitat_** | 0.20 (-) | Broad habitat and land use categories in 1000-meter area perimeter around survey area. Positive values (agriculture, water, island); negative (grassland, urban, forest) (Fig. S2) |
| **PC2_habitat_** | 0.77 (+) | Broad habitat and land use categories in 1000-meter area perimeter around survey area. Positive values (island, urban), negative (oak) (Fig. S2) |
| **Area of oak patch (m^2^)** | 0.78 (-) | Survey area plus area of extended perimeter that is oak habitat |
| **Survey area (m^2^)** | 0.30 (-) | Size of the total survey area |
| **Tree size** | 0.27 (+) | PC1 between DBH and tree height for trees in a survey period |

**Table S4**: Statistical significance for additive partitioning was assessed by comparing the proportion of null values that were greater (or less) then the estimate, given as a P-value using the software PARTITION (Veech and Crist, 2009). Significance values are bolded and underlined.

| **Level** | **P-value (S1)** | **P-value (S2)** | **P-value (S3)** |
| --- | --- | --- | --- |
| **α1 (within tree)** | **0** | **0** | **0** |
| **β1 (among trees)** | 1 | 1 | 1 |
| **α2 (within site)** | 1 | 1 | 1 |
| **β2 (among sites)** | **0.28** | 0.99 | 0.99 |
| **α3 (within region)** | 1 | 1 | 1 |
| **β3 (among region)** | **0** | **0** | **0** |

**Table S5**: Community composition (CA) gall morphotype loadings for site level CA (see Fig. 2).

| **Acronym** | **CA1** | **CA2** | **CA3** | **CA4** | **CA5** | **CA6** |
| --- | --- | --- | --- | --- | --- | --- |
| ***TT*** | -0.97 | 0.16 | 0.81 | -0.55 | -0.03 | -0.13 |
| ***AS*** | -0.92 | 0.12 | 0.63 | -0.30 | 0.00 | -0.28 |
| ***AKA*** | -0.85 | 0.44 | 0.12 | 0.02 | 0.08 | 0.19 |
| ***DP*** | -0.81 | 0.39 | -0.01 | -0.04 | 0.16 | 0.08 |
| ***ACO*** | -0.80 | 0.45 | -0.33 | -0.36 | -0.04 | -0.42 |
| ***AF*** | -0.73 | 0.63 | -0.31 | 0.46 | 0.46 | 0.74 |
| ***PB*** | -0.65 | 1.80 | -2.78 | -0.07 | 0.23 | -0.35 |
| ***ACG*** | -0.57 | 0.10 | 0.33 | 0.74 | 0.72 | 1.73 |
| ***XT*** | -0.54 | 0.03 | -0.09 | 0.10 | -0.33 | 0.05 |
| ***AG*** | -0.46 | -0.14 | 0.01 | 0.39 | 0.24 | -0.09 |
| ***AK*** | -0.33 | -0.21 | -0.09 | -0.02 | 0.21 | -0.05 |
| ***AP*** | -0.30 | -0.02 | -0.59 | 0.37 | -0.41 | 0.11 |
| ***ACH*** | -0.30 | -0.34 | 0.18 | 0.40 | -0.38 | 0.27 |
| ***AC*** | -0.27 | -0.37 | -0.10 | -0.26 | 0.00 | -0.08 |
| ***AQ*** | -0.27 | -0.29 | -0.10 | 0.32 | -0.09 | 0.07 |
| ***DME*** | 0.14 | 0.06 | 0.14 | 0.12 | -0.17 | 0.08 |
| ***DM.DS*** | 0.14 | -0.06 | -0.23 | -0.55 | -0.38 | 0.32 |
| ***DW*** | 0.18 | -0.85 | -0.21 | -0.09 | 0.29 | 0.15 |
| ***ACF*** | 0.22 | -1.02 | -0.30 | 0.16 | -0.25 | -0.44 |
| ***NW*** | 0.26 | 0.13 | 0.04 | 0.01 | 0.08 | -0.03 |
| ***BMI*** | 0.28 | -0.09 | 0.03 | 0.06 | 0.21 | -0.07 |
| ***AO*** | 0.40 | 0.22 | 0.04 | -0.03 | -0.05 | -0.06 |
| ***NSA*** | 0.66 | 0.27 | 0.12 | 0.01 | 0.02 | -0.02 |
| ***DC*** | 0.93 | -1.78 | -1.05 | -2.88 | 1.49 | 1.15 |

**Table S6**: Models (LMM, GLMM) with geographic + abiotic predictor variables with ΔAICc < 2.

| **Response variables** | **Predictor variables** | **K** | **AICc** | ΔAICc |
| --- | --- | --- | --- | --- |
| **CA1** | latitude | 5 | 97.92 | 0 |
| **CA1** | latitude + PC1_temp_ | 6 | 98.88 | 0.96 |
| **Gall abundance - NSA (log)** | latitude | 5 | 86.71 | 0 |
| **NSA abundance (log)** | latitude + PC1_temp_ + soil moisture | 7 | 148.91 | 0 |
| **Total gall abundance (log)** | latitude + PC1_temp_ | 6 | 38.55 | 0 |
| **Richness** | latitude + soil moisture+ PC1_temp_ + elevation + PC2_temp_ + PC1_precip_ | 9 | 261.16 | 0 |
| **Richness** | latitude + soil moisture | 5 | 262.73 | 1.57 |

**Table S7**: Models (LMM, GLMM) with geographic + habitat predictor variables with ΔAICc < 2.

| **Response** | **Predictor variables** | **K** | **AICc** | ΔAICc |
| --- | --- | --- | --- | --- |
| **CA1** | latitude | 4 | 105.17 | 0 |
| **Gall abundance – NSA (log)** | latitude | 4 | 90.12 | 0 |
| **NSA abundance (log)** | latitude | 4 | 166.6 | 0 |
| **NSA abundance (log)** | latitude + elevation | 5 | 167.31 | 0.72 |
| **Total gall abundance (log)** | latitude | 4 | 59.82 | 0 |
| **Total gall abundance (log)** | elevation | 4 | 60.01 | 0.19 |
| **Total gall abundance (log)** | latitude + elevation | 5 | 60.3 | 0.48 |
| **Richness** | latitude + elevation+ PC2_habitat_ + oak area | 6 | 268.76 | 0 |

**Table S8**: Models (LMM, GLMM) with predictor variables retained after model selection ΔAICc < 2 with abiotic and habitat variable separately.

| **Response** | **Predictor variables** | **K** | **AICc** | ΔAICc < 2. |
| --- | --- | --- | --- | --- |
| **CA1** | latitude | 5 | 97.92 | 0 |
| **CA1** | latitude + PC1_temp_ | 6 | 98.88 | 0.96 |
| **Gall abundance – NSA (log)** | latitude | 5 | 86.71 | 0 |
| **NSA abundance (log)** | latitude + PC1_temp_ | 5 | 164.82 | 0 |
| **NSA abundance (log)** | latitude | 4 | 166.6 | 1.78 |
| **Total gall abundance (log)** | latitude + PC1_temp_ | 5 | 45.01 | 0 |
| **Richness** | latitude + elevation+ PC1_temp_ + PC2_habitat_ + oak area | 7 | 264.97 | 0 |

**Table S9**: Within site abiotic and habitat factor CCA with presence/absence of cynipids on trees for each region. *P*-values reveal significant relationships. Boldened and underlined values < 0.05; bolded only < 0.10.

|  | R1 | R2 | R3 | R4 | R5 | R6 |
| --- | --- | --- | --- | --- | --- | --- |
| Soil moisture (corrected) | **0.003** | **0.042** | **0.001** | **0.001** | 0.929 | 0.625 |
| Soil temperature (corrected) | **0.02** | **0.001** | **0.024** | **0.08** | 0.514 | 0.19 |
| Distance to non-oak edge (m) | 0.513 | **0.074** | **0.014** | **0.001** | 0.933 | 0.32 |
| Tree Size (DBH, tree height) | 0.231 | 0.257 | 0.141 | **0.008** | 0.514 | **0.073** |
| Julian date | **0.001** | **0.001** | **0.016** | **0.001** | 0.859 | 0.662 |

**Figure S1**: Biplots showing PC1 and PC2 of a) temperature and b) precipitation BIOCLIM variables as they relate to study sites (see Table S1 for site acronyms).

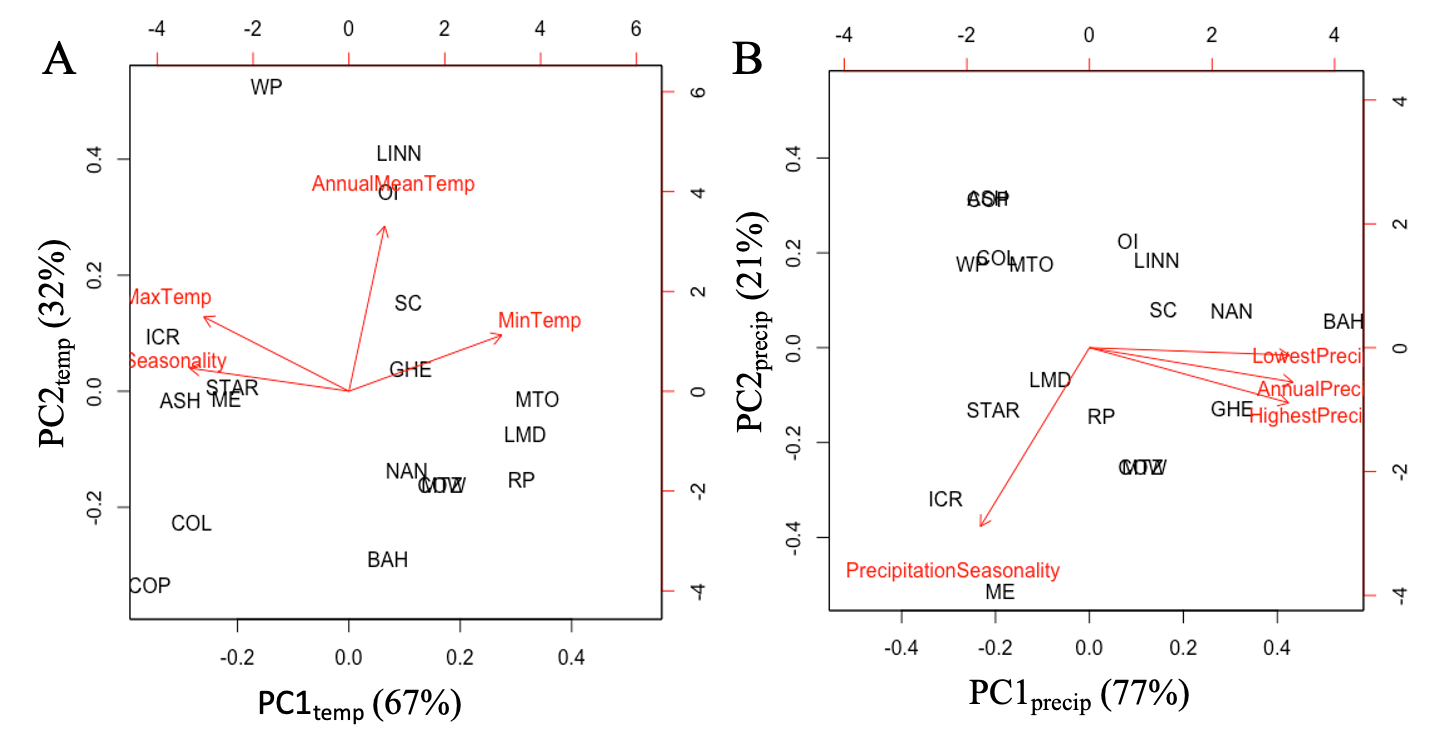

**Figure S2**: Biplot of PC1 and PC2 of broad habitat categories in 1-kilometer radius around site survey areas.

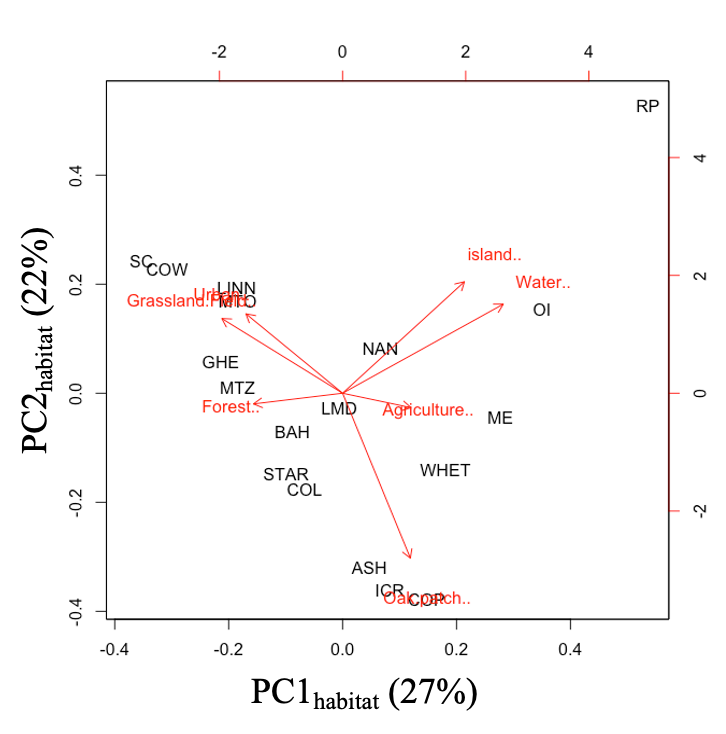

**Figure S3**: Additive partitioning of cynipid morphotype richness for each survey period. Richness was partitioned among (β-diversity) and within (α-diverity) regions, sites, and trees. Observed distributions are compared to null models that assumed a random distribution with 1000 randomizations using a Monte Carlo simulation based off data to estimate α and β values using the software PARTITION (Veech and Crist, 2009).

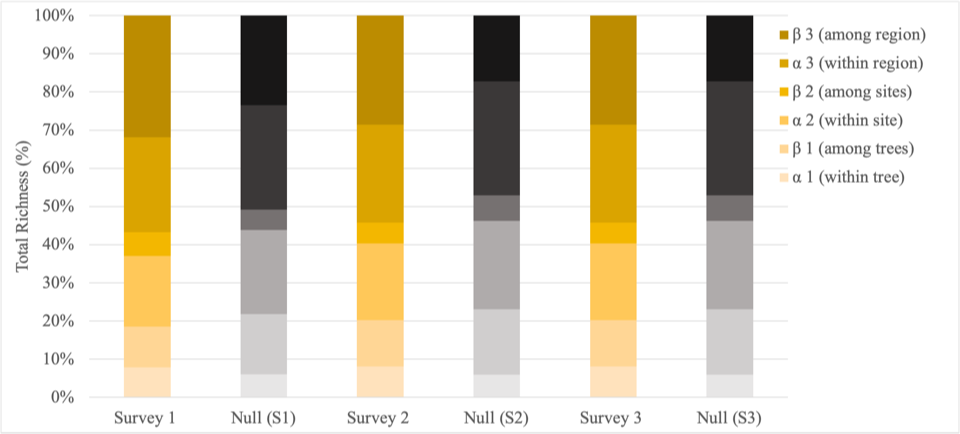

**Figure S4**: Species rarefaction curves, estimating cynipid morphotype richness in regions using surveyed trees as samples. All curves asymptote, suggesting sampling within regions was adequate.

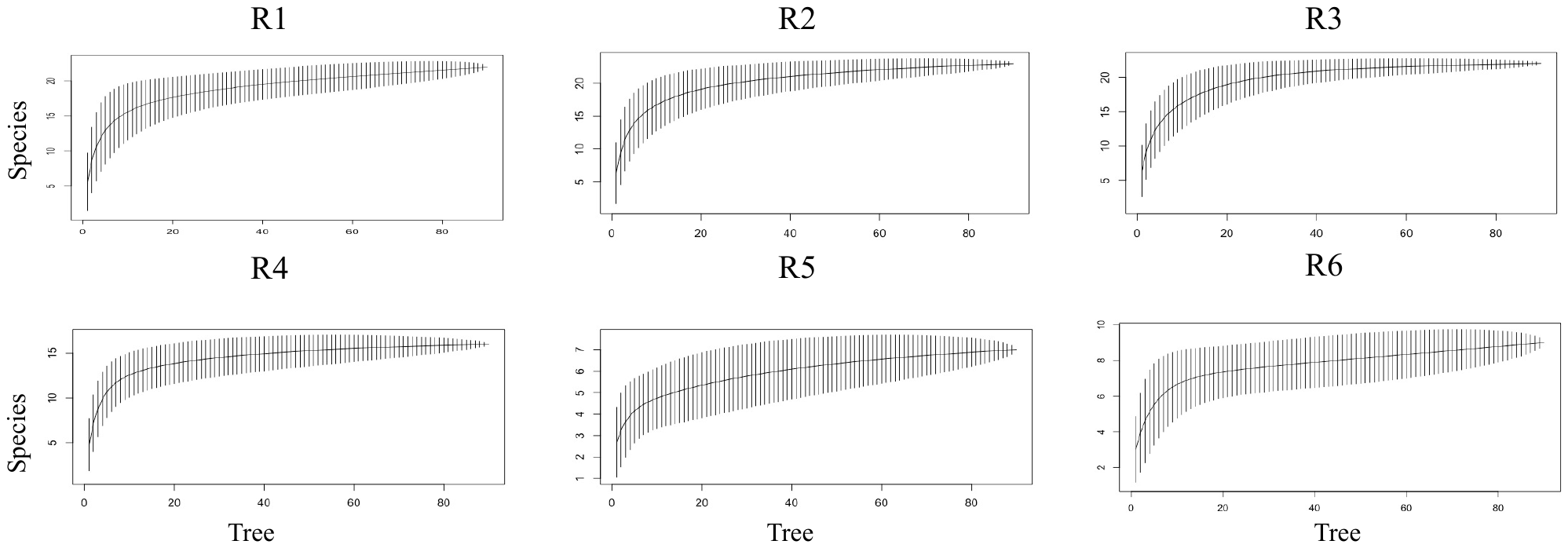

**Figure S5**: Total gall abundance (log-transformed) across latitude (log-transformed). Regions are colored coded. Solid black line represents regression line (LM) with 95% confidence intervals in dash green lines.

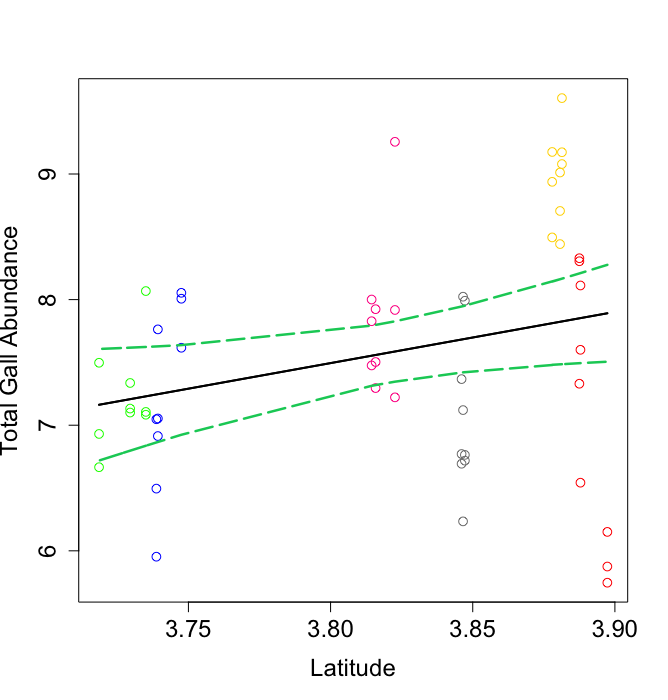

**Figure S6**: Abundance of *N. saltatorius* at sites in six regions. Regions and sites are colored coded (see Figure 1).

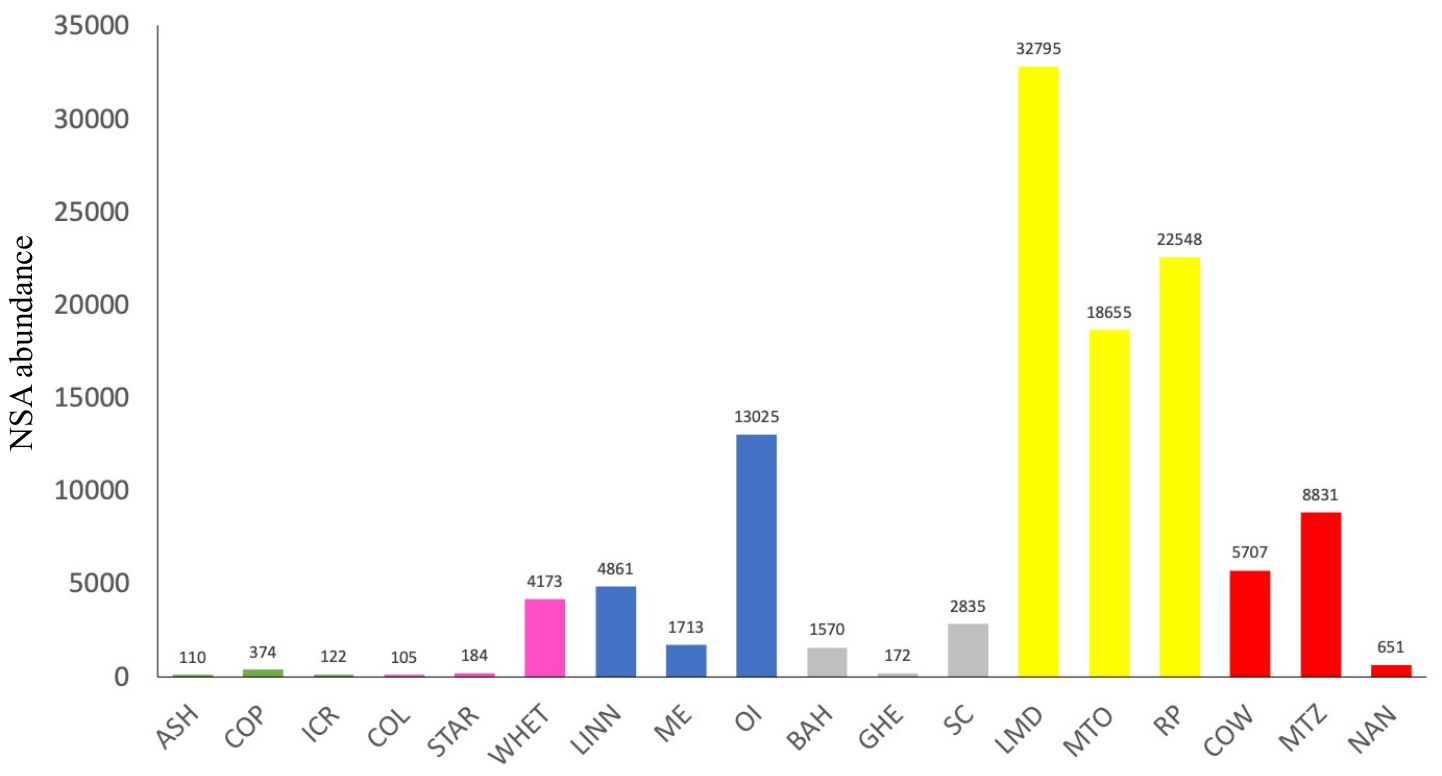

**Figure S7**: Biplots of CCAs with abiotic and habitat variables with presence/absence of cynipids on trees for each region. Open symbols represent trees, colored symbols morphotypes showing gall type groups (detachable, blue; integral, purple; stem, orange; focal species, black).

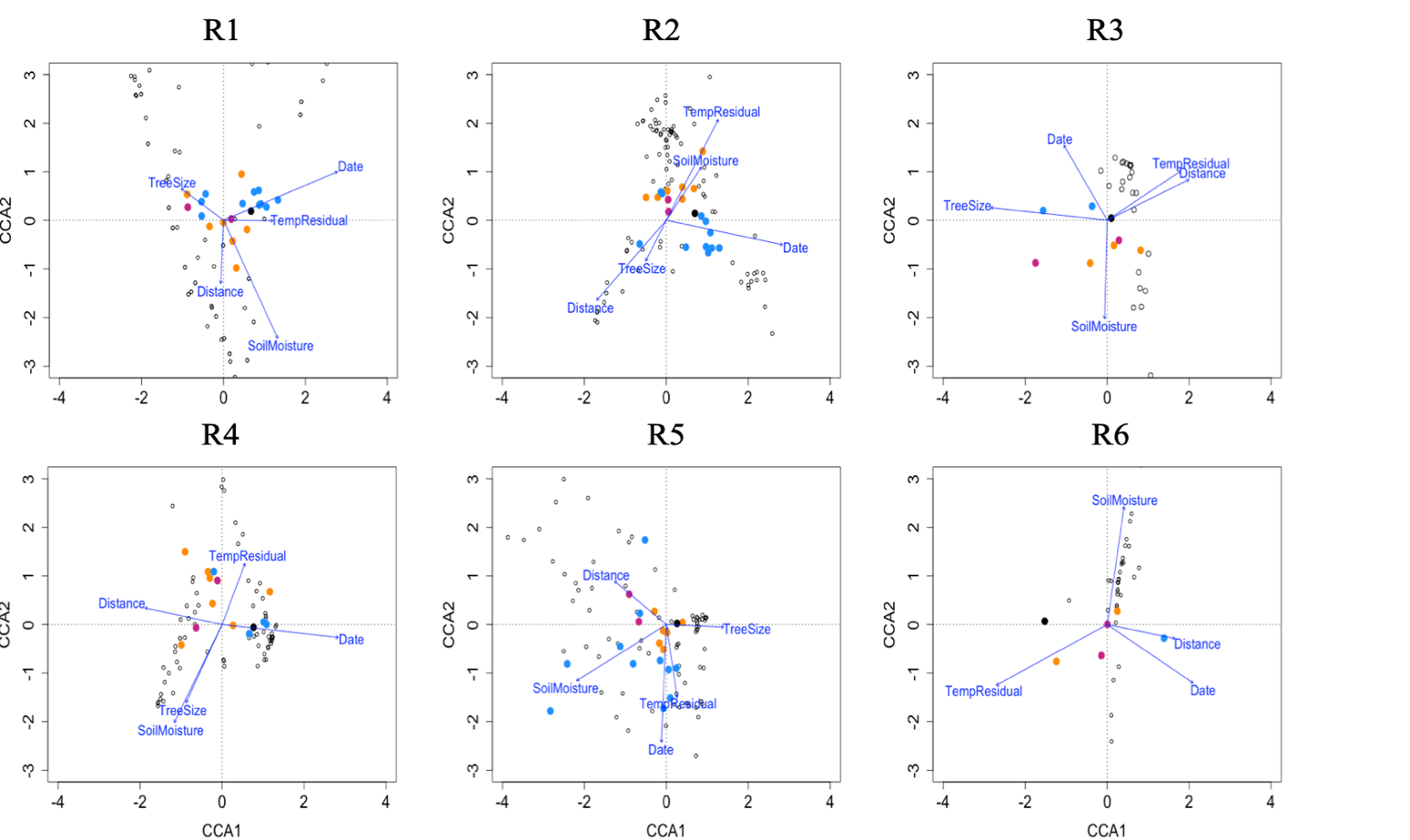
